# Double face of cytochrome c in cancers. New look into human breast ducts with Raman imaging

**DOI:** 10.1101/2021.05.25.445576

**Authors:** H. Abramczyk, B. Brozek-Pluska, M. Kopeć

## Abstract

Cytochrome c (Cyt c) is a key protein that is needed to maintain life (respiration) and cell death (apoptosis). The dual-function of Cyt c comes from its capability to act as mitochondrial redox carrier that transfers electrons between the membrane-embedded complexes III and IV and to serve as a cytoplasmic apoptosis-triggering agent, activating the caspase cascade.^1–6^ However, the precise roles of Cyt c in mitochondria, cytoplasm and extracellular matrix under normal and pathological conditions are not completely understood.^7–9^

To date, no pathway of Cyt c release that results in caspase activation has been compellingly demonstrated in any invertebrate.^10^ The significance of mitochondrial dysfunctionality has not been studied in ductal carcinoma to the best of our knowledge.^1^

Here we show that proper concentration of monounsaturated fatty acids, saturated fatty acids, cardiolipin and Cyt c is critical in the correct breast ductal functioning and constitutes an important parameter to assess breast epithelial cells integrity and homeostasis. We look inside human breast ducts answering fundamental questions about location and distribution of various biochemical components inside the lumen, epithelial cells of the duct and the extracellular matrix around the cancer duct during cancer development *in situ*. We found in histopathologically controlled breast cancer duct that Cyt c, cardiolipin, and palmitic acid are the main components inside the lumen of cancerous duct *in situ*. The presented results show direct evidence that Cyt c is released to the lumen from the epithelial cells in cancerous duct. In contrast the lumen in normal duct is empty and free of Cyt c. Our results demonstrate how Cyt c is likely to function in cancer development. We anticipate our results to be a starting point for more sophisticated *in vitro* and *in vivo* animal models. For example, the correlation between concentration of Cyt c and cancer grade could be tested in various types of cancer. Furthermore, Cyt c is a target of anti-cancer drug development ^11,12^ and a well-defined and quantitative Raman based assay for oxidative phosphorylation and apoptosis will be relevant for such developments.

## Introduction

Recent years have yielded exciting findings in the field of cancer cell metabolism, suggesting that change in the cellular redox status is important cancer driver, controlling various aspects of malignant progression. ^13^ Altered mitochondrial metabolism and redox state of cytochrome c (Cyt c) is being increasingly recognized as an important factor, triggering various processes in cancer development. ^5,6,13–17^ It has been more than two decades since the central role of Cyt c in mitochondrial pathway has been reported. ^1,5,6,14–17^ However, key issues regarding how Cyt c is released from mitochondria and from cells still remain largely unclear.^3,5,6,14–17^

Cyt c belongs to family of heme containing metalloproteins. Cyt c is located in the intermembrane space of mitochondria and released into bloodstream during pathological conditions. Circulating in blood Cyt c level is suggested to be a novel *in vivo* marker of mitochondrial injury after resuscitation from heart failure and chemotherapy.^18^ Various existing techniques such as enzyme-linked immunosorbent assays (ELISA), Western blot, high performance liquid chromatography (HPLC), spectrophotometry and flow cytometry have been used to estimate Cyt c concentration. However, the implementation of these techniques at POC (point of care) application is limited due to longer analysis time, expensive instruments and expertise needed for operation.^18^ Moreover, none of the methods used to control Cyt c concentration can provide direct evidence about the role of cytochrome c in apoptosis and oxidative phosphorylation, because they are not able to monitor the amount of cytochrome in specific organelles such as mitochondria, cytoplasm, or extracellular matrix.

Here we show that Raman spectroscopy and Raman imaging are a promising label-free methods to estimate not only the level of Cyt c in fast analysis of clinical practice, but also to identify localization and biochemical content in epithelial cells of the duct and in the extracellular matrix.

Until now, no technology has proven effective for detecting Cyt c concentration in specific cell organelles. Therefore, existing analytical technologies cannot detect the full extent of Cyt c localization inside and outside specific organelles. In Raman imaging we do not need to disrupt cells to break open the cells and release the cellular structures to learn about their biochemical composition.

Cyt c is not only serving as an cell death biomarker (apoptosis), but is also a key protein that is needed to maintain life (respiration). Thus, it is of great importance to understand the role of Cyt c in certain diseases at cellular level. ^6^ Here we will concentrate on breast cancer.

Here we show that mitochondrial content of Cyt c is critical in the correct breast ductal functioning and constitutes an important parameter to assess breast epithelial cells integrity and homeostasis. We look inside human breast ducts answering fundamental questions about location and distribution of various biochemical components inside the lumen, epithelial cells of the duct and the extracellular matrix around the cancer duct during cancer development *in situ*.

We studied oncogenic processes that characterize human breast cancer (ductal cancer *in situ* (DCIS) and infiltrating ductal carcinoma (IDC)) based on the quantification of cytochrome redox status by exploiting the resonance-enhancement effect of Raman scattering.

In this paper we explore a hypothesis involving the role of reduction-oxidation pathways related to Cyt c in cancer development. Here we show that Raman spectroscopy and Raman imaging are competitive clinical diagnostics tools for cancer diseases linked to mitochondrial dysfunction and are a prerequisite for successful pharmacotherapy of cancer.

## 1. Results

To properly address redox state changes of mitochondrial cytochromes in breast cancers by Raman spectroscopy and imaging, we systematically investigated how the Raman method responds to *in vitro* human cells and *ex vivo* human tissues. *In vitro* experiments will allow to study a single cell by reducing the role of cell-to cell interactions. The *ex vivo* human tissue experiments will extend our knowledge on the influence of environment on cancer development.

Figure 1 shows the cross section through the normal breast duct obtained by Raman imaging. Details of the experimental method used to create Raman image are given in Supplementary Materials. The Raman image is compared with the microscopy image. The characteristic vibrational spectra for different areas of the breast tissue are also presented in Fig.1. One can see from Fig.1. that there is an almost perfect match between the morphological features and Raman images. However, Raman imaging provides additional information, which is not available from histology, microscopy, mammography, and fluorescence. It is biochemical information. To understand biochemical information that is provided from Raman images we need to associate the characteristic features with the breast morphology. Briefly, the normal organization of ducts demonstrates lumen surrounded by epithelial cells aligned in a polar manner so their apical side faces the lumen. These cells are surrounded by the basement membrane. The next layers represents extracellular matrix consisting of fibroblasts and the stroma, which is predominantly, but not exclusively, composed of connective tissue and adipose tissue. Schematic structures of epithelial tissue, stromal and adipocyte cells around the normal breast duct are presented in Scheme 1.

**Figure 1.**
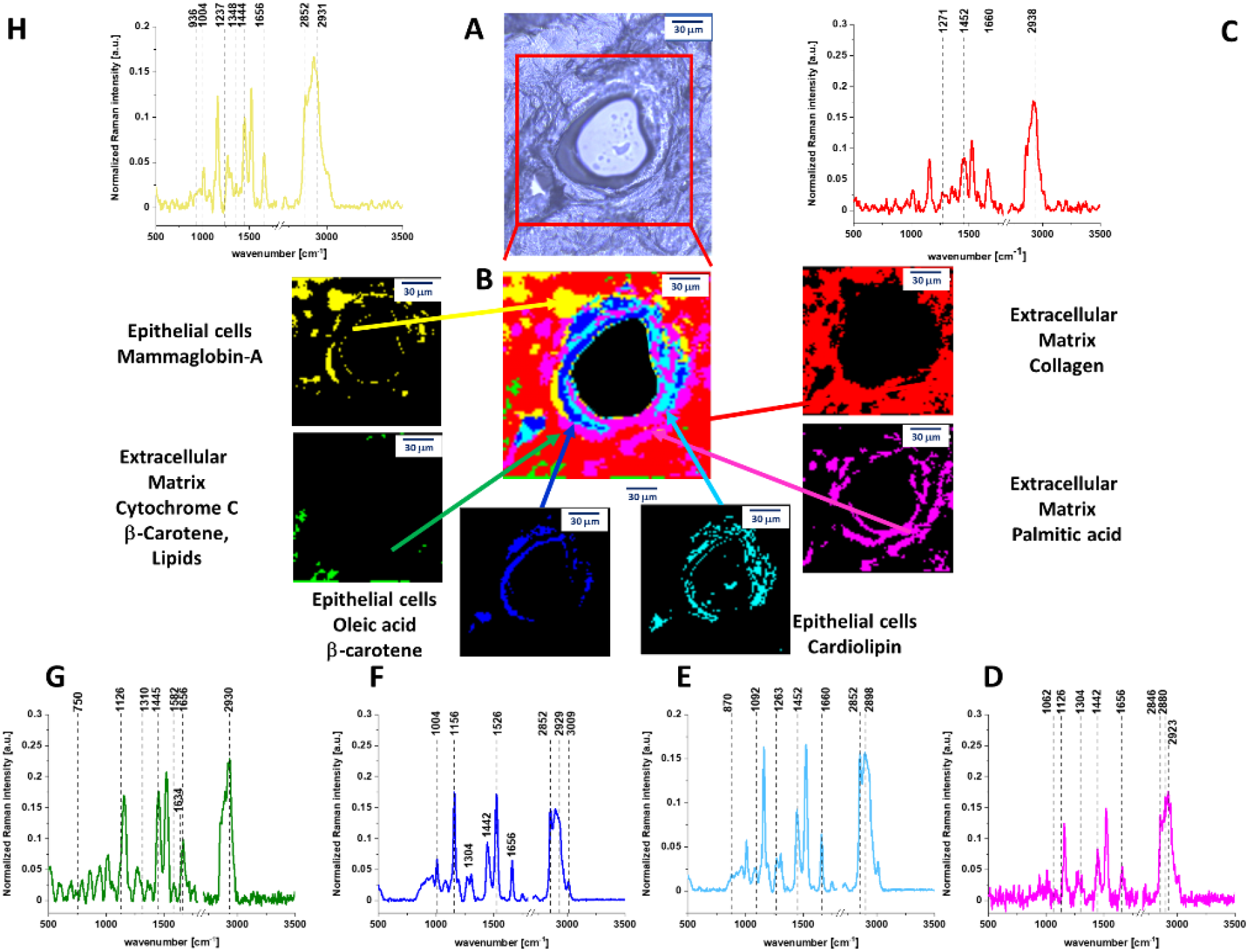
(A) Microscopy image of human normal duct, (B) Raman imaging of human normal duct obtained by using Cluster Analysis and Raman images and average Raman spectra of all clusters identified by Cluster Analysis: (C) collagen (red), (D) palmitic acid (pink), (E) cardiolipin (turquoise), (F) oleic acid, (G) cytochrome c, (H) mammaglobin-A (yellow).

**Scheme 1.**
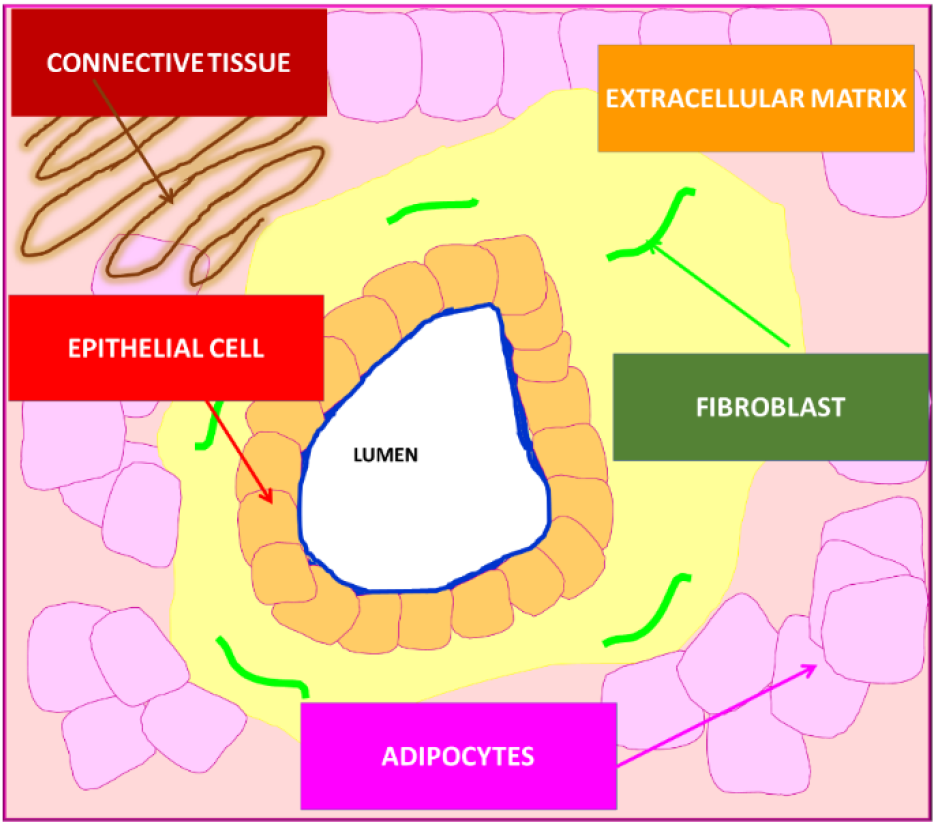
Schematic representation of the structure of human normal duct.

Comparing Raman and microscope images from Fig. 1 with the Scheme 1 it is easy to identify all morphological features of the normal duct.

One can see that yellow-blue line around the black duct represents normal epithelial cells that are lined along the intact basement membrane and do not proliferate inside the lumen and outside through the basement membrane. Raman images provides information not only about morphology, but also about biochemistry of these structures that is given by the Raman spectra in Fig.1.

The Raman spectra presented in Fig.1 show the biochemical composition of the structures. One can see from Fig. 1B that the lumen is empty (black colour) with no Raman spectra indicating that there are no epithelial/mesenchymal cells inside the normal duct. The epithelial cells of the normal duct contain oleic acid, β-carotene, cardiolipin, palmitic acid. The epithelial cells are dominated by monounsaturated oleic acid derivatives composed of glyceryl trioleate and carotenoids (Fig.1F). Indeed, the characteristic Raman vibrations of carotenoids with resonance peaks at 1156 and 1526 cm^−1^ are clearly visible in Fig.1F. The peaks at 2852, 2928, 3009 cm^−1^ correspond to the vibrations of monounsaturated oleic acid ^19–21^. To show the perfect match between Raman spectra of carotenoids and monounsaturated oleic acid derivatives in human normal duct and Raman spectra of pure isolated compounds the correlation analysis was performed (Pearson correlation coefficient was equal 1.0 at the confidence level 0.95), see Supplementary Materials and Pearson correlation coefficient for all components.

The extracellular matrix (red colour in Fig.1C.) is dominated by collagen (see Supplementary Materials). Small concentration of oxidized cytochrome c (Fe^3+^ green colour in Fig.1.) was found with a characteristic Raman peaks at 750, 1126, 1582 cm^−1^ and 1634 cm^−1^.^22^ The oxidized form of cytochrome c (Fe^3+^) can induce caspase activation in the process of apoptosis, while the reduced form (Fe^2+^) cannot.^23^ The oxidized cytochrome c (Fe^3+^) is not bound to cardiolipin and can participate in electron shuttling of the respiratory chain and in oxidative phosphorylation.^24^ Cardiolipin, is abundantly present in mitochondria in the inner mitochondrial membrane, where it constitutes about 20% of the total lipid composition. ^17^

The band at 1656 cm^−1^ represents Amide I vibration of proteins and C=C stretching vibration of unsaturated lipids (mainly monounsaturated oleic acid).^19,22^

In contrast to the normal duct in Fig.1. the cancerous duct presented in Fig.2. shows that the normal organization of the epithelial cells is lost and the lumen is filled with the cancerous cells. It would be extremely interesting to learn what chemical substances are released from the epithelial cells into the lumen during cancer development, because monitoring biochemical alterations would drive the progress on mechanisms of cancer to limits just unimaginable a few years ago. To our best knowledge this a first report about the chemical composition of cancerous and normal human breast ducts.

**Figure 2.**
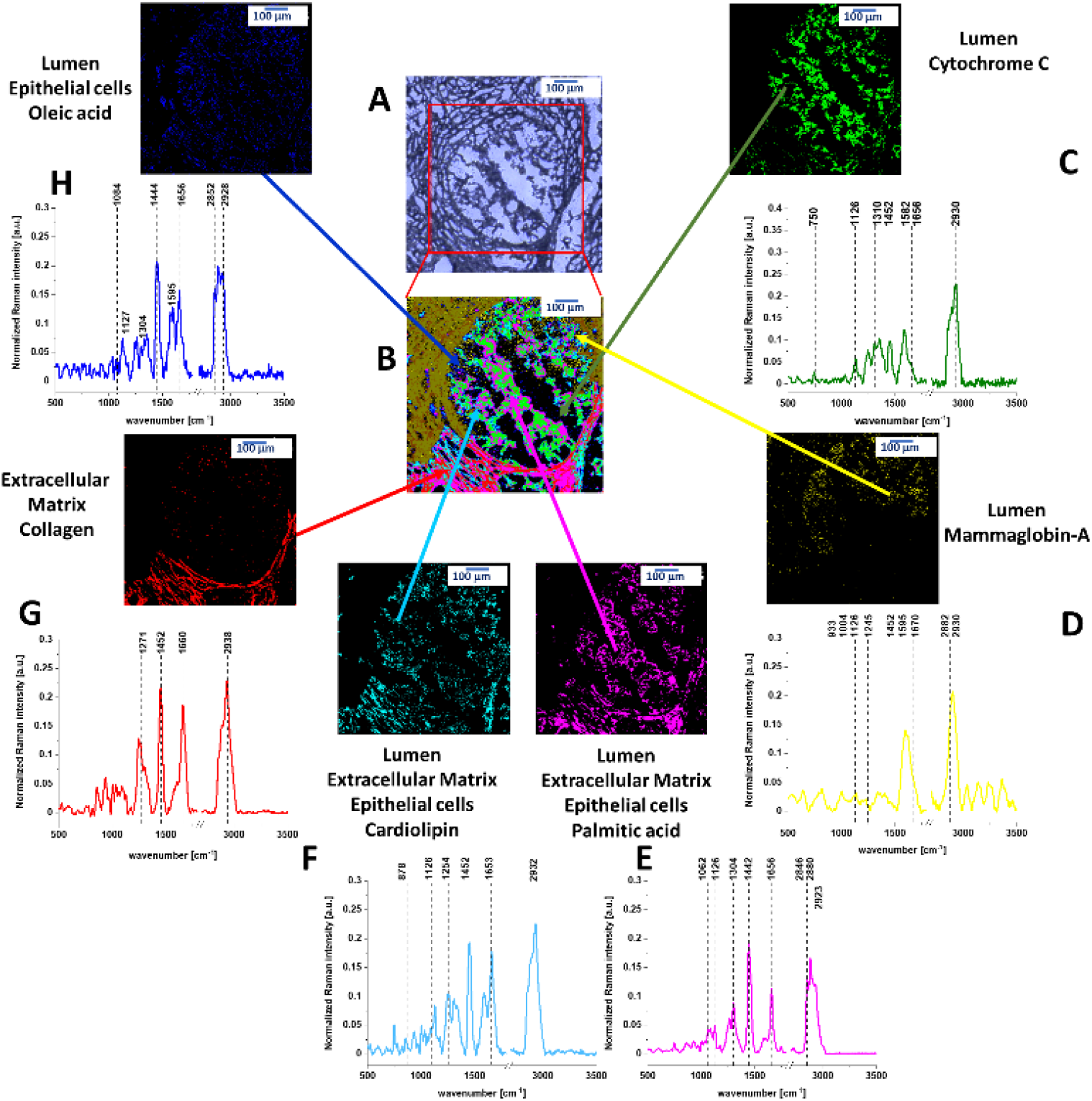
(A) Microscopy image of human cancerous duct, (B) Raman imaging of human cancerous duct obtained by using Cluster Analysis and Raman images and average Raman spectra of all clusters identified by Cluster Analysis: (C) cytochrome c (green), (D) mammaglobin-A (yellow), (E) palmitic acid (pink), (F) cardiolipin (turquoise), (G) collagen (red), (H) oleic acid (blue).

Fig. 2 shows the Raman image of the cross section through the cancerous duct. Fig. 2 demonstrates that the biochemical profile of the lumen of the cancerous duct contains four main components: cytochrome c (green colour), cardiolipin (turqoise), palmitic acid (pink colour) and mammaglobin-A (yellow colour), (Details in Supplementary Materials) in contrast to the lumen in the normal duct which is empty (Fig. 1). The cytochrome c represents the reduced form (Fe^2+^) as the Raman signal at 1582 cm^−1^ of the reduced form is an order higher than that of the oxidized form (Fe^3+^) (see Supplementary Materials). The Raman spectrum of Cyt c in Fig. 2 shows that Cyt c is bound to a lipid identified as a cardiolipin (1452 cm^−1^). The band at 1452 cm^−1^ of cardiolipin does not overlap with the C-H deformation bands of saturated lipids of palmitic acid at 1442 cm^−1^. In addition, one can see that the Raman intensity of the band at 1656 cm^−1^ in Fig. 2C corresponding to C=C vibrations of unsaturated lipids is much lower than that of reduced cytochrome c at 1582 cm^−1^ in contrast to the normal duct (Fig. 1G). This is a result of decrease of C=C vibration from monounsaturated oleic acid, which is evident from comparison Raman signals at 1656 cm-1 between Fig. 1F and 2H.

Cyt c is mostly protonated meaning that most Cyt c bonds via electrostatic bonds to acidic phospholipids, particularly cardiolipin. Cardiolipin-bound Cyt c, probably does not participate in electron shuttling of the respiratory chain.^24^ It indicates that the process of oxidative phosphorylation (respiration) becomes less effective in cancer cells (known as Wartburg effect).

On the other hand, the reduced form of cytochrome c (Fe^2+^) cannot induce caspase activation and the process of apoptosis in cancerous cells becomes less efficient.^23^

The lipid profile of the lumen in the cancerous breast duct in Fig. 2 is dominated by a mixture of cardiolipin (turquoise colour), palmitic acid (pink colour) and oleic acid (blue colour) with no presence of carotenoids in contrast to normal epithelial cells in the normal duct filled with monounsaturated oleic acid derivatives composed of glyceryl trioleate and carotenoids (Fig.1). We found that the ratio between the monounsaturated oleic acid and palmitic acid is 3:1 in normal duct and 1:1 in cancerous duct. We found that the ratio between cytochrome c in normal duct and in cancerous duct is 1:47. We found also that the ratio between cardiolipin in normal duct and in cancerous duct is 1:1.6. Details are given in Supplementary Materials.

The alterations in the lipid composition in the epithelial cells must have very serious consequences. Incorporation of saturated lipids, such as cardiolipin and palmitic acid into lipid membranes is known to stiffen a membrane. Such membranes can be described as “a rigid amorphous glass state ^24^ leading to distortions and deformations. The extracellular matrix around the cancerous duct is dominated by a network consisting of complementary regions of collagen (red colour) and other proteins (mammaglo-bin-A, yellow colour). Strong fluorescence at 599 nm (yellow dark colour in the upper left corner in Fig. 2B) for the excitation at 532 nm is also observed.

First we will concentrate on contribution of cytochrome c to the cancerous duct. Fig. 1 shows that in normal duct, Cyt c is located around the epithelial cells in the extracellular matrix. Fig.2 shows that in the cancerous duct Cyt c is located in the lumen. To show the perfect match between Raman spectra in the lumen of the human cancerous duct and Raman spectrum of isolated Cyt c the correlation analysis was performed (Pearson correlation coefficient was equal 1.0 at the confidence level 0.95), see Supplementary Materials.

Therefore, the results in Fig. 2 provides the first direct evidence for the release of Cyt c from epithelial cells into the lumen of the cancerous duct *in situ* where it has a reduced form. The mechanisms how Cyt c is released to the lumen of the cancerous duct is still unknown, but in the view of the presented results they must be related to the lipid composition of the epithelial cells.

At normal physiological conditions, Cyt c is located in the mitochondrial intermembrane/intercristae spaces of cells, where it functions as an electron shuttle in the respiratory chain and interacts with cardiolipin.^17^ It is commonly believed that proapoptotic oncogenic stimuli induce the permeabilization of the outer membrane allowing for Cyt c release to cytosol. In the cytosol, Cyt c mediates the allosteric activation of apoptosis-protease activating factor 1, which is required for the proteolytic maturation of caspase-9 and caspase-3. Activated caspases ultimately lead to apoptotic cell dismantling.^17^ There are a few possible scenarios of the outer membrane permeabilization such as induction of mitochondrial permeability transition, Bcl-family proteins and mitochondrial outer membrane permeabilization, Volume-dependent, MPT-independent mechanisms of cytochrome c release, caspase-2-mediated release of Cyt c.^5^

Our results suggest that permeability of the membranes is simply related to the modifications of their biochemical composition of lipids during cancer development. We found that the protein/lipid profiles inside the lumen and in the epithelial cells of the cancerous duct are markedly different for the normal and the cancerous ducts. Our results (Fig.1) demonstrates that the epithelial cells of the normal duct are dominated by monounsaturated fatty acids that contributes to a proper membrane fluidity of the epithelial cells. The cardiolipin and palmitic acid are located around the epithelial cells in the extracellular matrix. In contrast, the epithelial cancerous cells and the lumen are enriched with saturated lipids (cardiolipin and palmitic acid). Membranes of cells are the primary target for injury and their damage and are highly dependent on their physical properties and lipid organization. The abnormal proportion between saturated and unsaturated fatty acids that we found in the epithelial cells of the cancerous duct (Fig.2) effects fluidity of the membranes leading to distortions and deformations and decrease of mechanical stability.^24–28^ Membrane fluidity is a key property for maintaining cell functionality, and depends on lipid composition and cell environment.^28^ The effects of mechanical deformations due to modifications of fluidity result in expelling cardiolipin, Cyt c and palmitic acid into the lumen of the cancerous duct. The distortion of the epithelial cells allows Cyt c to be released to lumen.

In the view of the results presented so far one can propose discrete sequence of biochemical events that lead to malignant transformation of the epithelial cells in the normal breast duct. First, the upregulation of glycolysis ^29–34^ leads to enhanced synthesis of fatty acids *de novo*.^35–41^ *De novo* fatty acids synthesis changes biochemical composition of the epithelial cells as one can see from the comparison of Figures 1 and 2. The altered fluidity of the membranes of epithelial cells leads to mechanical deformations allowing for Cyt c release to lumen of the duct. In this hypothesis the release of Cyt c is a result, not a cause, of malignant transformation due to lipid reprogramming in cancer development.^22,42,43^ To date, no pathway of Cyt c release that results in caspase activation has been compellingly demonstrated in any invertebrate, despite the presence of homologues of many of the molecules that mediate and/or regulate the intrinsic pathway of apoptosis in vertebrate cells. Furthermore, little is known about the extent of Cyt c release (if any) in cells that do not die.^6^

We anticipate our results to be a starting point for more sophisticated analysis of statistical significance. That is why we tested the correlation between concentration of Cyt c and cancer grade in ductal breast cancer, which has not been studied in the literature to the best of our knowledge.

Fig. 3 shows Raman intensity of the characteristic vibration of cytochrome c and b (750, 1126, 1337 and 1584 cm^−1^) as a function of grade for human breast normal (G0) and cancer human tissues (G1, G2, G3).

**Figure 3.**
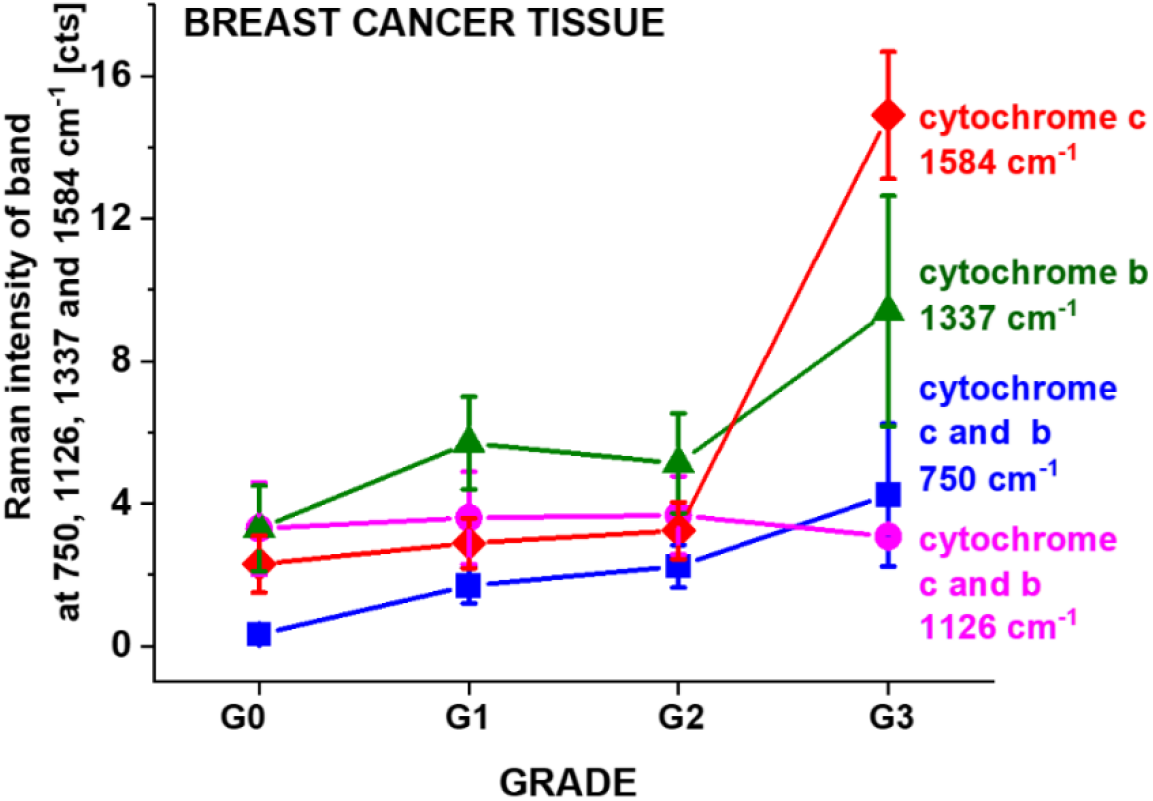
Raman intensities of the bands at 750, 1126, 1337 and 1584 cm^−1^ as a function of grade for breast normal (G0) and cancer (invasive ductal cancer) human tissue (G1, G2, G3) (average +/− SD, number of patients n=39). Detailed statistical analysis (one-way ANOVA) has been shown in Figures 4 and 5.

**Figure 4.**
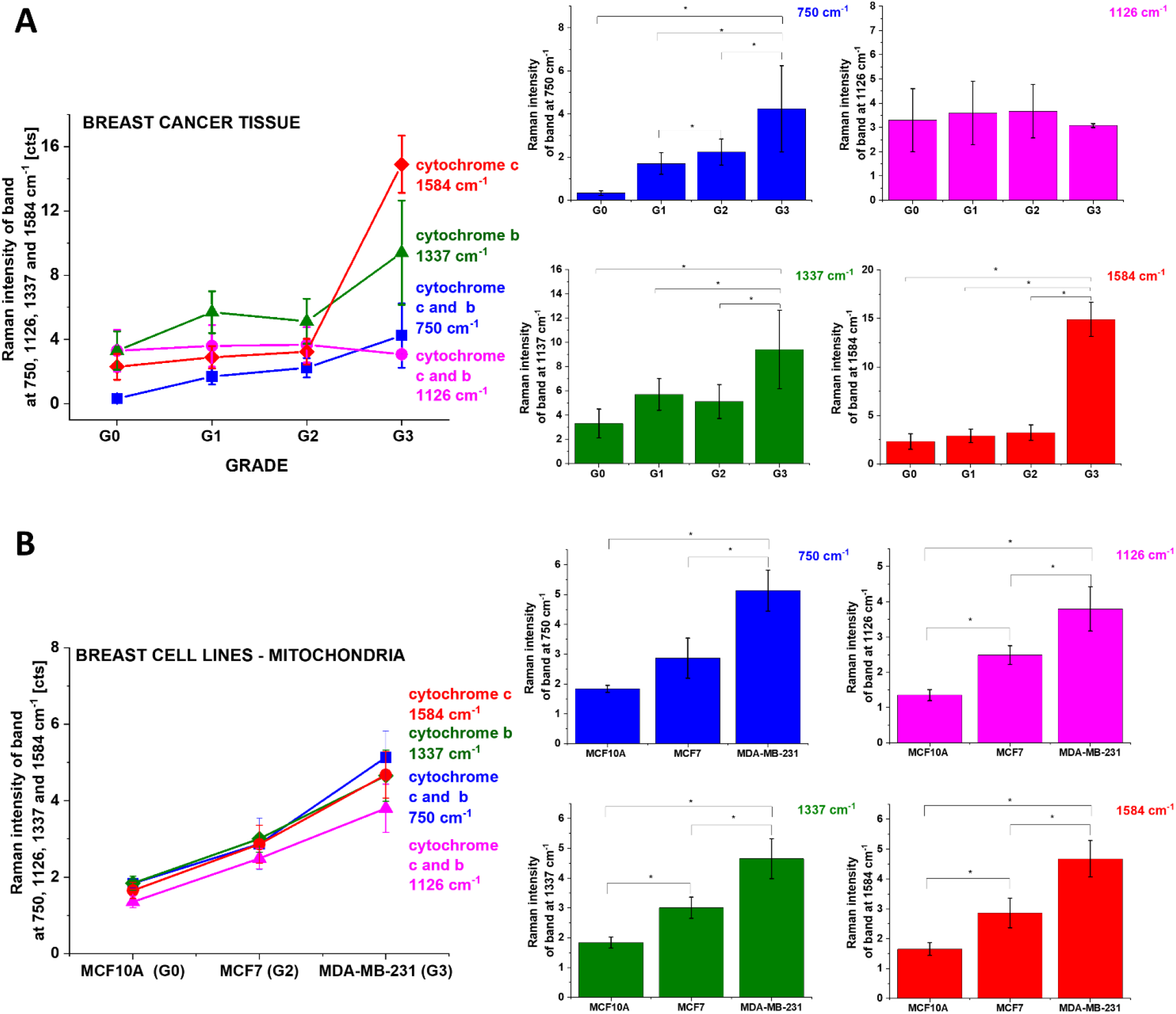
The Raman intensities I_750_, I_1126_, I_1337_, I_1584_ of Cyt c and Cyt b bands in human breast tissues (A) and in mitochondria of single breast cells *in vitro* (B) as a function of breast cancer grade malignancy G0-G3 at excitation 532 nm, average+/− SD, the statistically significant result have been marked with asterisk. The one-way ANOVA using the Tukey test was used to calculate the value significance, asterisk * denotes that the differences are statistically significant, *p*-values ≤ 0.05 were accepted as statistically significant.

**Figure 5.**
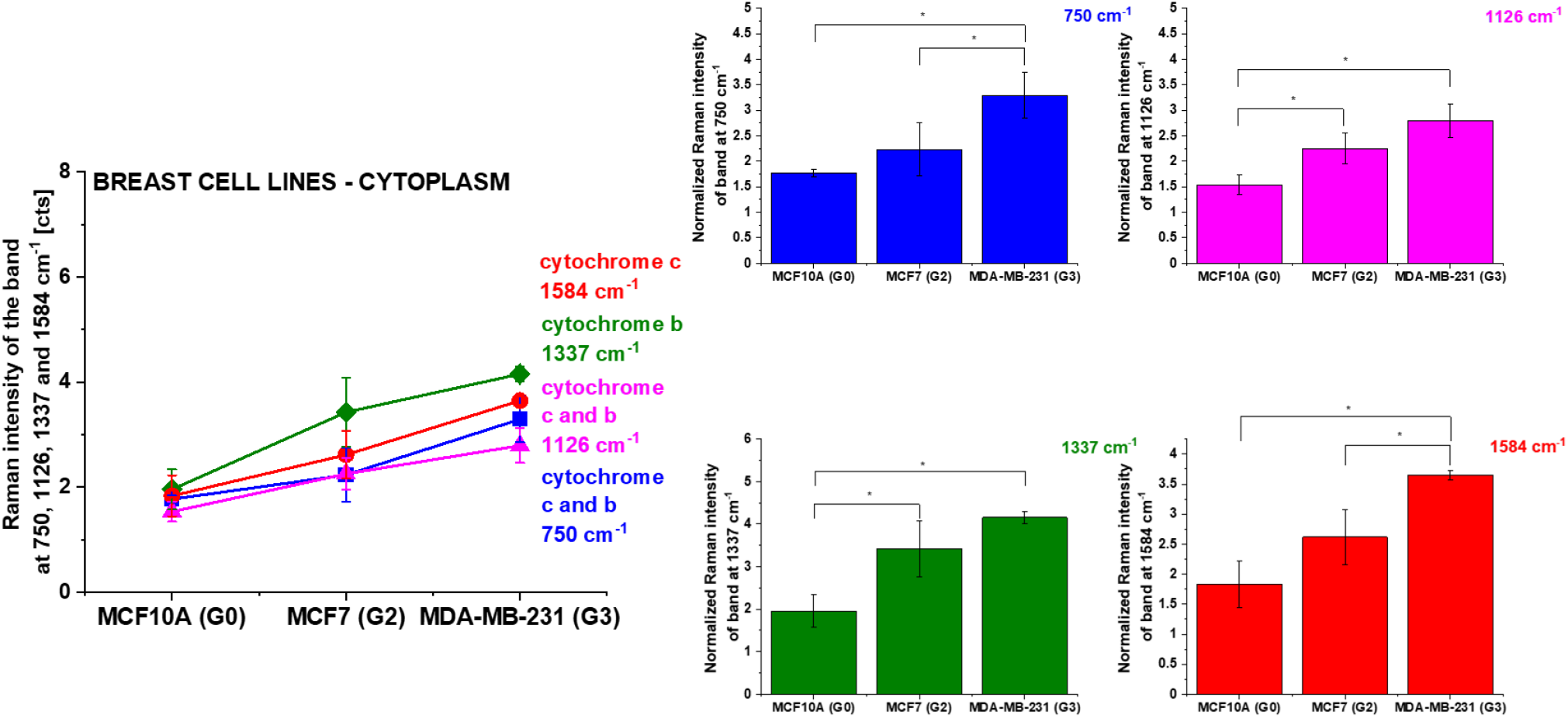
The Raman intensities of Cyt c and Cyt b in cytoplasm of single breast cells *in vitro*: I_750_, I_1126_, I_1337_, I_1584_ as a function of breast cancer grade malignancy G0-G3 at excitation 532 nm, the statistically significant result have been marked with asterisk. The one-way ANOVA using the Tukey test was used to calculate the value significance, asterisk * denotes that the differences are statistically significant, *p*-values ≤ 0.05 were accepted as statistically significant.

The total number of patients n was 39, for each patient thousands spectra (typically 6400) obtained from Raman imaging were used for averaging. Based on the average values obtained for the Raman biomarkers of Cyt c and b we obtained a plot as a function of breast grade malignancy. One can see from Fig. 3 that the global concentration of reduced cytochrome c in the breast tissue (reflected by the Raman intensity of the band at 1584 cm^−1^) increases with cancer aggressiveness. The correlation between Cyt c concentration and cancer aggressiveness is characterized by gradually increasing Raman signal at 1584 cm^−1^ indicating progressive redox state changes.

To understand the results presented in Fig.3 we need to examine normal and cancerous cells *in vitro*, where we will be able to directly monitor the concentration of cytochrome in separate organelles of the epithelial cells: mitochondria, cytosol, lipid droplets and nucleus by using Raman imaging.

Let us concentrate on Cyt c concentration in mitochondria in breast cells as a function of cancer aggressiveness. We studied human breast cells of normal cells (MCF 10A), slightly malignant cells (MCF7) and highly aggressive cells (MDA-MB-231) by means of Raman microspectroscopy at 532 nm. We visualized localization of cytochromes by Raman imaging in the major organelles of cells. We demonstrated that the “redox state Raman marker” of the ferric low spin heme in Cyt c at 1584 cm^−1^ as well as mixed vibrations (cytochrome c and b) can serve as a sensitive indicators of cancer aggressiveness. We compared concentration of Cyt c vs. grade of cancer aggressiveness in cancer tissues and single cells in specific organelles in cells: nucleous, mitochondria, lipid droplets, cytoplasm, and membrane. (Cluster Analysis results for human breast cell are presented in Supplementary Materials).

Fig. 4 shows the Raman intensities I_750_, I_1126_, I_1337_, I_1584_ cm^−1^ of vibrational peaks of Cyt c and b in human breast tissues and in mitochondria of single breast cells as a function of breast cancer aggressiveness.

One can see from Fig. 4 that the intensity of Raman biomarker at 1584 cm^−1^ corresponding to concentration of Cyt c in mitochondria of a single cell increases with breast cancer aggressiveness. The intensity of Raman biomarker at 1337 cm^−1^ corresponding to concentration of Cyt b also increases with breast cancer aggressiveness. The same tendency can be seen for intensities of Raman biomarkers at 750 cm^−1^ and 1126 cm^−1^ corresponding to concentration of Cyt c and b. Moreover, Fig. 4 demonstrates that both breast cancer tissue and breast cancer cell lines *in vitro* show similar trends. The higher concentration of Cyt c in mitochondria of cancer cells (MCF7 (G2) and MDA-MB-231 (G3)) *in vitro* when compared with the normal cells (MCF10A (G0)) as presented in Fig. 4 indicates that the concentration of Cyt c is upregulated in breast cancer cells.

The results show that the concentration of Cyt c increases, not decreases with cancer aggressiveness. It indicates that the netto concentration of Cyt c in mitochondria is higher than release to cytoplasm (if any). This finding reflects the dual face of Cyt c: apoptosis and oxidative phosphorylation. The balance between cancer cells proliferation (oxidative phosphorylation) and death (apoptosis) decide about level of cancer development.

The Cyt c concentration in mitochondria as a function of cancer aggressiveness reflects its contribution to oxidative phosphorylation and apoptosis. Normal cells (G0) primarily produce energy through glycolysis followed by mitochondrial citric acid cycle and oxidative phosphorylation. The signal at 1584 cm^−1^ for normal cells (G0 in Fig. 4B) represents predominantly the oxidized form (Fe^3+^) of Cyt c unbound to cardiolipin. It indicates that both apoptosis can be induced and electron shuttling between the complex III, Cyt c, and complex IV can occur leading to effective oxidative phosphorylation (respiration). In contrast, for cancerous cells (G2, G3) the concentration of reduced form (Fe^2+^) of Cyt c bound to cardiolipin increases. Cardiolipin-bound Cyt c, probably does not participate in electron shuttling of the respiratory chain^28^, and reduced cytochrome cannot induce caspase and apoptosis process.

Our results show also that cancer cells predominantly produce their energy through the oxidative phosphorylation, in contrast to Warburg hypothesis that most cancer cells produce their energy through a high rate of glycolysis followed by lactic acid fermentation even in the presence of abundant oxygen. However, the effectiveness of the oxidative phosphorylation decreases with cancer aggressiveness due to higher concentration of cardiolipin bound to Cyt c.

To better understand mechanism of competition between apoptosis and oxidative phosphorylation we monitored Cyt c concentration in cytoplasm. In normal cells Cyt c is found in the mitochondria. The release of Cyt c into the cytoplasm is believed to induce the non-inflammatory process of apoptosis.^5,17^ When it is transferred to the extracellular space, it can cause inflammation. The assessment of Cyt c in the extracellular space might be used as a biomarker for determine mitochondrial damage and cell death.

To estimate the release to cytoplasm we monitored the Cyt c concentration in cytoplasm of breast cancer cells compared to the normal cells. We studied concentration of Cyt c and b using Raman markers I_750_, I_1126_, I_1337_, I_1584_ in cytoplasm as a function of cancer aggressiveness. Figure 5 shows Raman intensities of Cyt c and Cyt b in cytoplasm of single *in vitro* cells: I_750_, I_1126_, I_1337_, I_1584_ as a function of breast cancer malignancy at excitation 532 nm.

Detailed inspection into Fig.5 shows that the concentration of Cyt c in cytoplasm increases in cancerous cells compared to normal one. We have performed the One Way ANOVA analysis of the data presented in Figure 5 and at the 0.95 confidence level, the statistically significant result have been marked with asterisk.

The level of 1.83 for G0 and 3.64 for G3 for Raman intensity corresponds to the Cyt c concentration of 22 μM (based on Figure S6A in Supplementary Materials) and 82 μM (based on Figure S6C), respectively (see Supplementary Materials).

It indicates that we observe the additional release of Cyt c to cytoplasm with increasing cancer aggressiveness when compared with the normal cells (G0).

The same tendency as for the concentration of Cyt c in cytoplasm of single cells *in vitro* presented in Fig. 5 was observed for the global concentration of Cyt c with cancer aggressiveness in breast cancer tissue (Fig.3 and 4). The levels of 2.3 for G0 and 14.9 for G3 for Raman intensity correspond to the Cyt c concentrations of 19 μM (based on Figure S6B) and 266 μM (based on Figure S6D), respectively (see Supplementary Materials).

The higher differences between concentration of Cyt c in normal (G0) and cancerous (G3) tissues results from the extracellular matrix contribution. In the tissue the combined effect of localization in mitochondria, the release of cytochrome c from mitochondria into the cytoplasm and from cytoplasm into extracellular space is monitored. The results of this paper indicate that the release of Cyt c to extracellular space is the critical mechanism in the process of cancer development.

To summarize, the results provide evidence on the prominent role of cytochrome c in the intrinsic pathway of apoptosis and oxidative phosphorylation vs cancer aggressiveness. Nevertheless, there is still much to learn. Here we demonstrate that Raman imaging reveal new expanses on the role of Cyt c in cancer biology and mechanisms od apoptosis and oxidative phosphorylation. We anticipate our results to be a starting point for more sophisticated *ex vivo* human tissues, *in vitro* and *in vivo* animal models. For example, the correlation between concentration of Cyt c and cancer grade could be tested in various types of cancer. Furthermore, Cyt c is a target of anti-cancer drug development ^10–12^ and a well-defined and quantitative Raman based assay for oxidative phosphorylation and apoptosis will be relevant for such developments.

## Materials and Methods

### Reagents

All reagents were purchased from Sigma Aldrich (Poland) unless otherwise stated. Cytochrome c (no. C2506), cardiolipin (no. C0653).

### Ethics statement

All the conducted studies were approved by the local Bioethical Committee at the Polish Mother’s Memorial Hospital Research Institute in Lodz (53/216) and by the institutional Bioethical Committee at the Medical University of Lodz, Poland (RNN/323/17/KE/17/10/2017). Written consents from patients or from legal guardians of patients were obtained. All the experiments were carried out in accordance with Good Clinical Practice and with the ethical principles of the Declaration of Helsinki. Spectroscopic analysis did not affect the scope of surgery and course and type of undertaken hospital treatment.

### Patients

In the presented studies the total number of patients diagnosed with breast cancer was 39. All patients were diagnosed with ductal carcinoma in situ (DCIS) *in situ* or invasive ductal carcinoma and treated at the M. Copernicus Voivodeship Multi-Specialist Center for Oncology and Traumatology in Lodz.

### Tissues samples collection and preparation for Raman spectroscopy

Tissue samples were obtained during routine surgery. The fresh, non-fixed samples were used to prepare 16 micrometers sections placed on CaF_2_ substrate for Raman analysis. The adjacent slices were used for histopathological analysis, which was performed by professional pathologists from Medical University of Lodz, Department of Pathology, Chair of Oncology. The types and grades of cancers were diagnosed according to the criteria of the Current WHO by pathologists from Medical University of Lodz, Department of Pathology, Chair of Oncology.

### Subculture of Cell Lines

A human breast MCF10A cell line (CRL10317, ATCC) was grown with completed growth medium: MEGM Kit (Lonza CC3150) without gentamycin-amphotericin B mix (GA1000) and with 100 ng/ml cholera toxin; a slightly malignant human breast MCF7 cell line (HTB22, ATCC) in Eagle’s Minimum Essential Medium (ATCC 30-2003) with 10% fetal bovine serum (ATCC 30-2020) and highly aggressive human breast MDA-MB-231 cell line (HTB26, ATCC) in Leibovitz’s L15 Medium (ATCC 30-2008) with 10% fetal bovine serum (ATCC 30-2020). All human breast cell lines were maintained at 37°C in a humidified atmosphere containing 5% CO_2_. Cells were seeded on CaF_2_ window in 35mm Petri dish at a density of 5×10^4^ cells per Petri dish the day before examination. Before Raman examination, cells were fixed with 4% formalin solution (neutrally buffered) and kept in phosphate buffered saline (PBS, Gibco no. 10010023) during the experiment.

### Raman imaging for assessing cytochrome *c* release in human tissues and *in vitro* cells

The status of Cyt c (whether intact in the mitochondria or released) was examined by WITec (Ulm, Germany) alpha 300 RSA+ confocal microscope by recording Raman spectra and images. All images were acquired by the experimental set up consisting of diode laser 532 nm, the fibre of 50 μm, a monochromator Acton-SP-2300i and a CCD camera Andor Newton DU970-UVB-353. The excitation line was focused on the sample through a 40x dry objective (Nikon, objective type CFI Plan Fluor C ELWD DIC-M, numerical aperture (NA) of 0.60 and a 3.6–2.8 mm working distance) for tissue measurements and 40x water dipping objective (Zeiss W plan-Apochromat, VIS-IR, N numerical aperture (NA) of 1.0 and a 2.5 mm working distance) for human breast cell lines. The average laser excitation power was 10 mW for 532, with an integration time of 1.0 sec for low frequency range and 0.5 for high frequency range. An edge filters were used to remove the Rayleigh scattered light. A piezoelectric table was used to record Raman images. The cosmic rays were removed from each Raman spectrum (model: filter size: 2, dynamic factor: 10) and the smoothing procedure: Savitzky–Golay method was also implemented (model: order: 4, derivative: 0). Data acquisition and processing were performed using WITec Project Plus software.

### Cluster analysis

Spectroscopic data were analyzed using Cluster Analysis method. Briefly Cluster Analysis is a form of exploratory data analysis in which observations are divided into different groups that have some common characteristics – vibrational features in our case. Cluster Analysis constructs groups (or classes or clusters) based on the principle that: within a group the observations must be as similar as possible, while observations belonging to different groups must be different.

The partition of n observations (x) into k (k≤n) clusters S should be done to minimize the variance (Var) according to the formula:

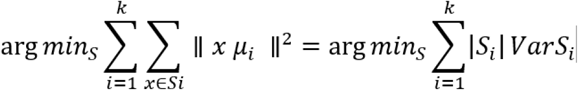

where *μ_i_* is the mean of points *S_i_*.

Raman maps presented in the manuscript were constructed based on principles of Cluster Analysis described above. Number of clusters was 6 (the minimum number of clusters characterized by different average Raman spectra, which describe the variety of the inhomogeneous biological sample). The colors of the clusters correspond to the colors of the average Raman spectra of collagen (red), Cyt c (green), oleic acid and β-carotene (blue), palmitic acid (pink), mammaglobin-A (yellow), cardiolipin (turquoise).

#### Basis analysis

The Basis analysis was performed based on the Raman spectra recorded for pure collagen, cytochrome c, oleic acid and β-carotene, palmitic acid, mammaglobin-A, and cardiolipin. During the analysis each measured spectrum of the 2D spectral array of the analyzed human breast sample was compared to the spectra of pure chemical components mentioned above using a least square to fit each convergence to minimize the fitting error D described by equation:

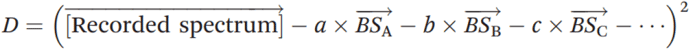

by varying the weighting factors a, b, c,… of the basis spectra 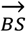.

**Table 1.**
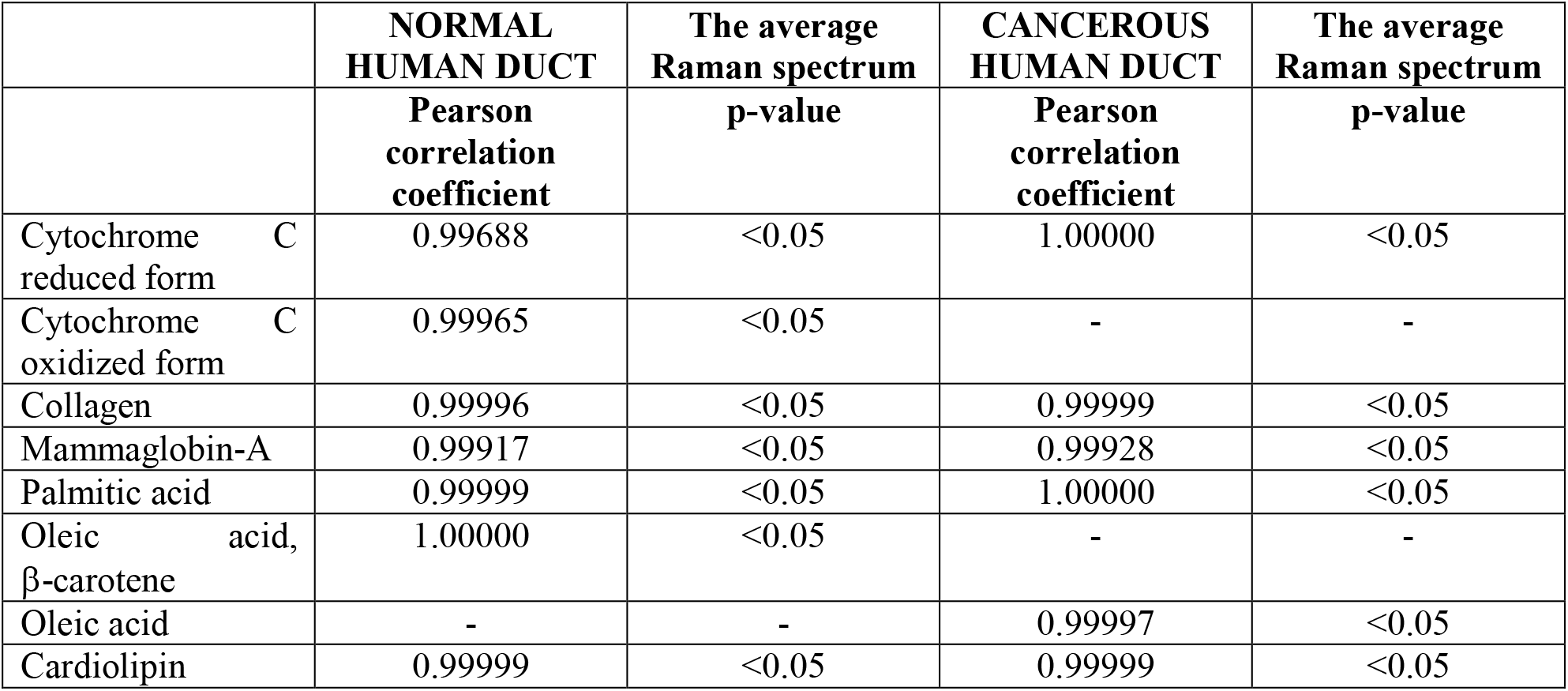
Pearson correlation coefficients obtained for comparison of the average Raman spectra typical for normal and cancerous human breast duct and the Raman spectra characteristic for pure components such as: oleic acid, β-carotene, palmitic acid, mammaglobin-A, collagen, cytochrome c, cardiolipin.

## Acknowledgements

This work was supported by NCN Poland grant UMO-2019/33/B/ST4/01961.

## SUPPLEMENTARY MATERIALS

Figure S1. shows the microscopy image, the Raman image, the histopathological image, the comparison of average Raman spectra obtained by Cluster Analysis Method and the Raman spectra characteristic for pure chemical components: oleic acid, β-carotene, palmitic acid, mammaglobin-A, collagen, cytochrome c, cardiolipin for normal human breast duct.

**Figure S1.**
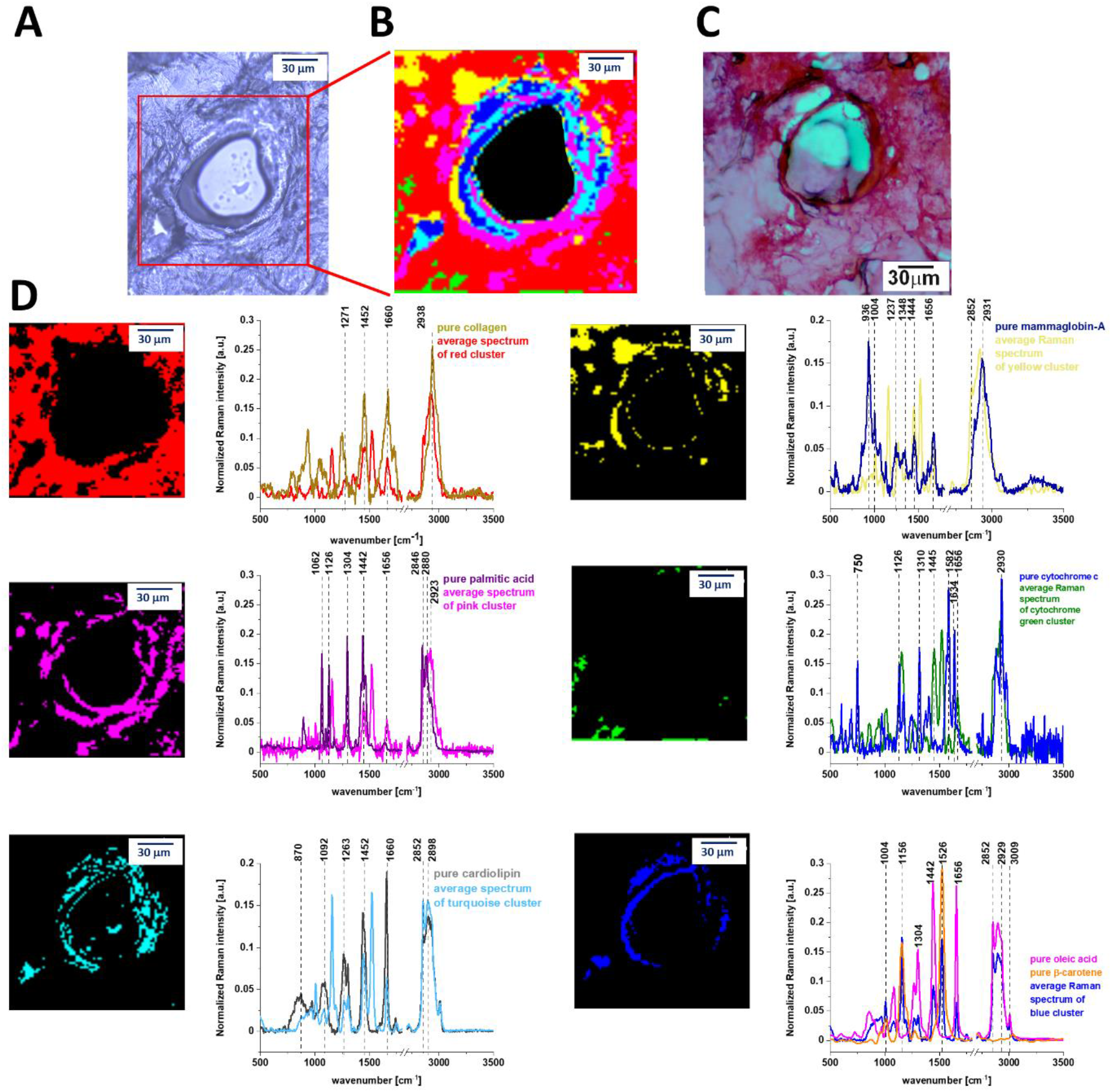
(A) Microscopy image of human normal breast duct, (B) the Raman image of human normal breast duct obtained by Cluster Analysis, (C) the histopathological image of human normal breast duct and (D) the comparison of average Raman spectra obtained by Cluster Analysis Method and the Raman spectra characteristic for pure chemical components: oleic acid, β-carotene, palmitic acid, mammaglobin-A, collagen, cytochrome c, cardiolipin.

Figure S2. the microscopy image, the Raman image, the histopathological image, the comparison of average Raman spectra obtained by Cluster Analysis Method and the Ramana spectra characteristic for pure chemical components: oleic acid, β-carotene, palmitic acid, mammaglobin-A, collagen, cytochrome c, cardiolipin for cancerous human breast duct.

**Figure S2.**
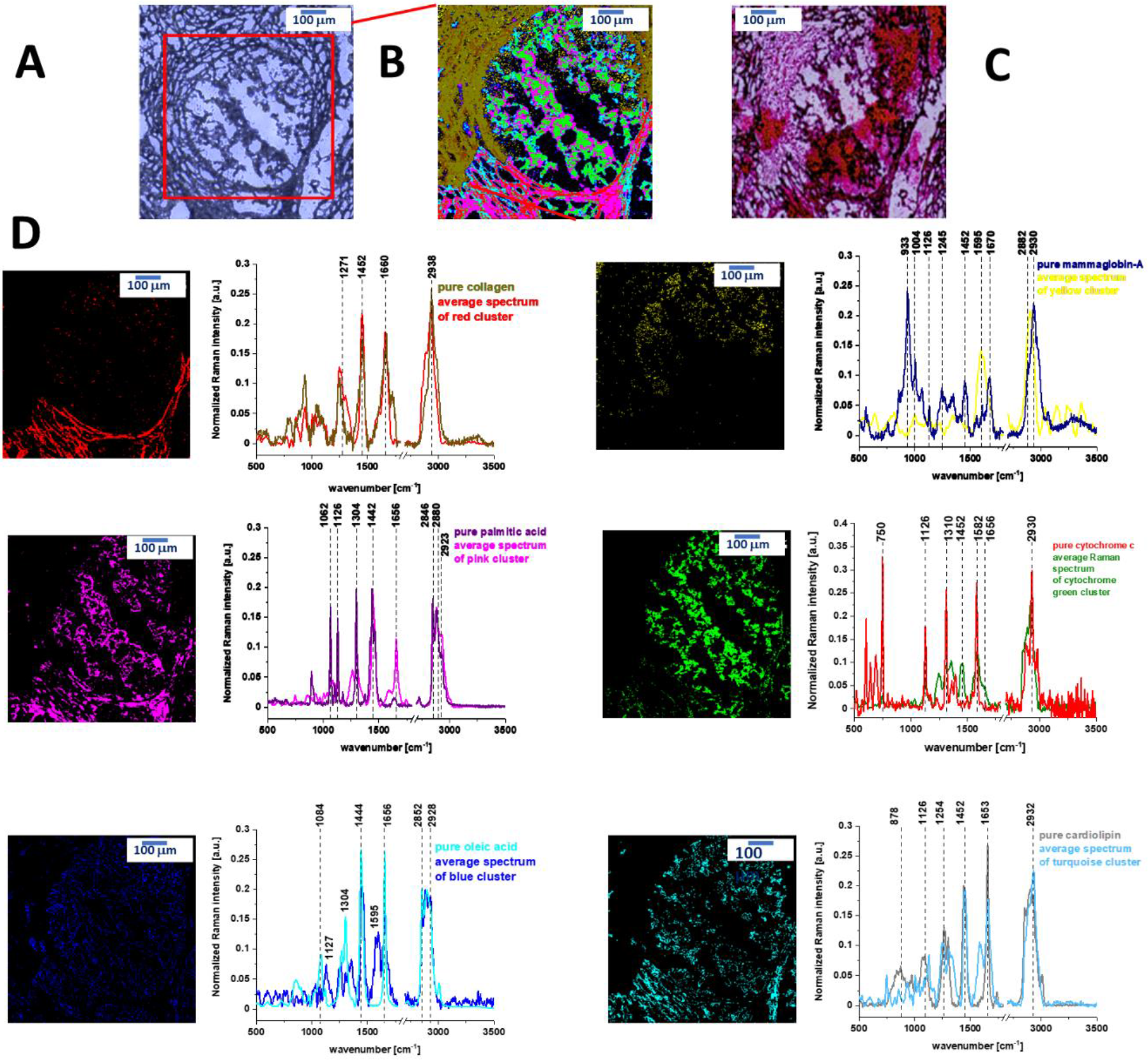
(A) Microscopy image of human cancerous breast duct, (B) the Raman image of human cancerous breast duct obtained by Cluster Analysis, (C) the histopathological image of human cancerous breast duct and (D) the comparison of average Raman spectra obtained by Cluster Analysis Method and the Raman spectra characteristic for pure chemical components: oleic acid, β-carotene, palmitic acid, mammaglobin-A, collagen, cytochrome c, cardiolipin.

Figure S3. shows the results of Basis analysis performed for normal human breast duct based on the Raman spectra of pure: oleic acid, β-carotene, palmitic acid, mammaglobin-A, collagen, cytochrome c, cardiolipin.

**Figure S3.**
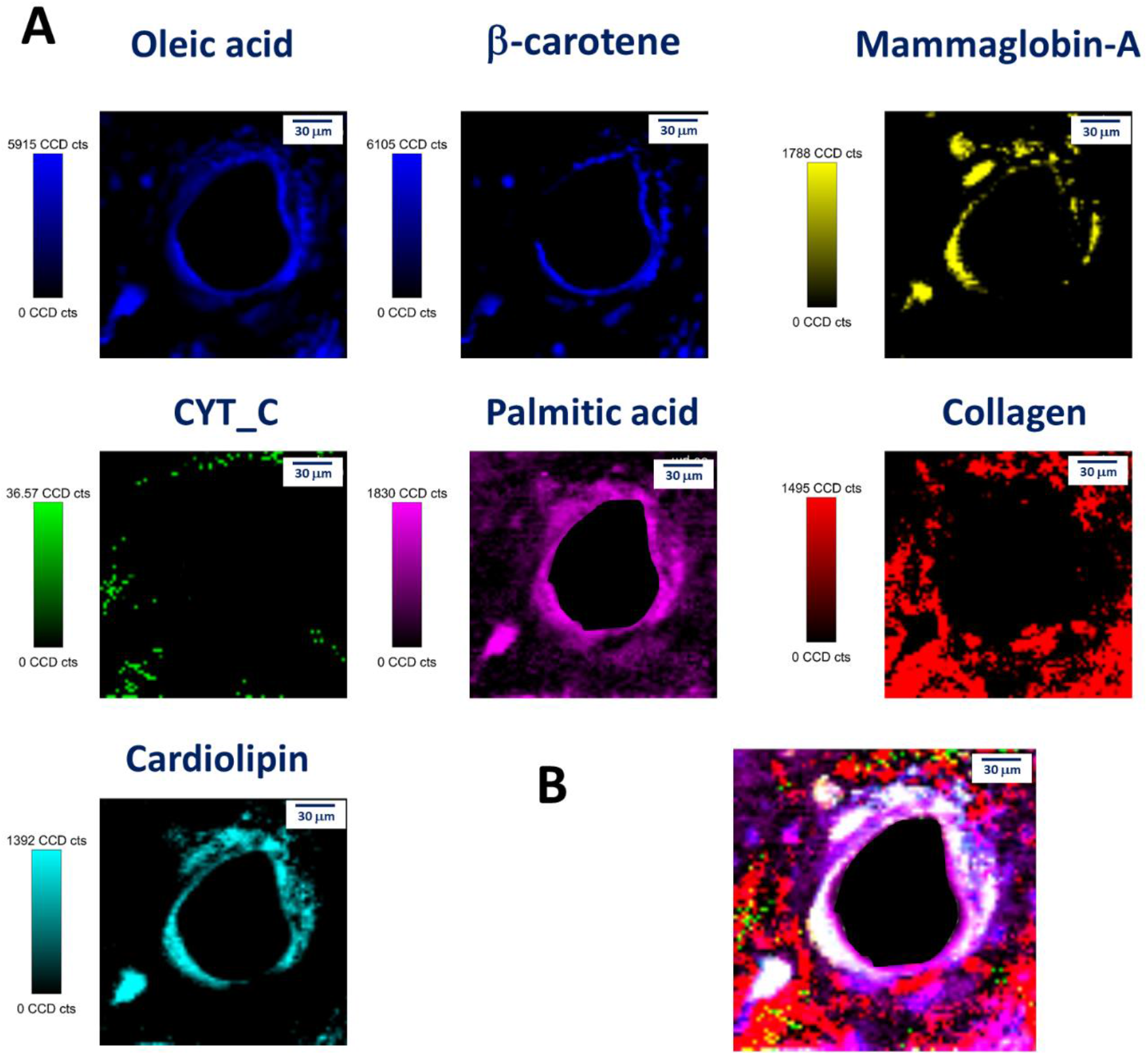
(A) Basis analysis performed for normal human breast duct based on the Raman spectra of pure: oleic acid, β-carotene, palmitic acid, mammaglobin-A, collagen, cytochrome c, cardiolipin; the images obtained for single chemical components and (B) the combination of images obtained for single chemical components shown on panel (A).

Figure S4. shows the results of Basis analysis performed for cancerous human breast duct based on the Raman spectra of pure: oleic acid, palmitic acid, mammaglobin-A, collagen, cytochrome C, cardiolipin.

**Figure S4.**
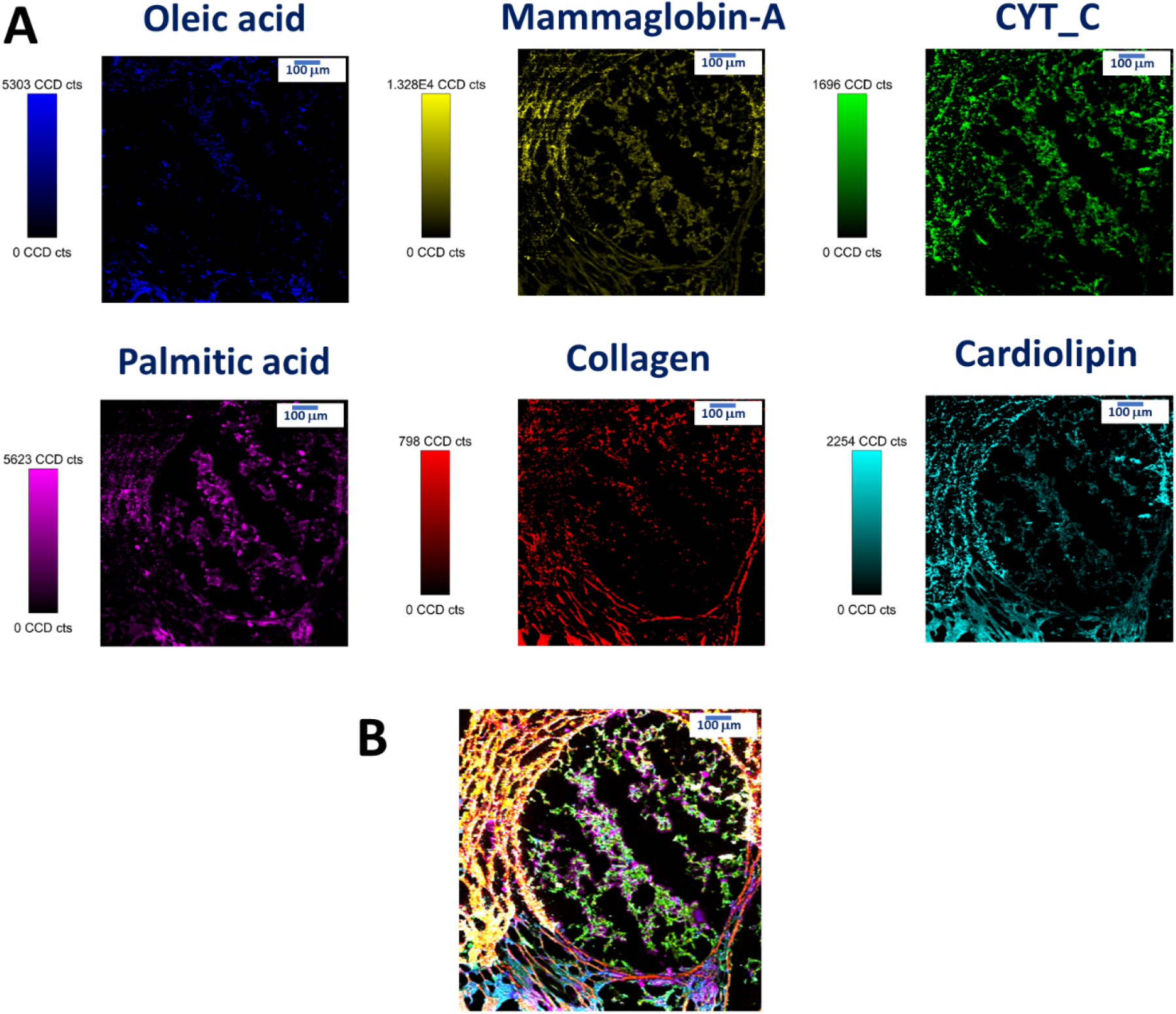
(A) Basis analysis performed for cancerous human breast duct based on the Raman spectra of pure: oleic acid, palmitic acid, mammaglobin-A, collagen, cytochrome c, cardiolipin; the images obtained for single chemical components and (B) the combination of images obtained for single chemical components shown on panel (A).

Figure S5 shows the microscopy image (A), the Raman image of a single cell MDA-MB-231 of highly aggressive breast cancer obtained by Cluster Analysis (B) and corresponding average Raman spectra of different organelles of the cell: nucleus (red), cell membrane (light grey), lipid structures (blue and orange), cytoplasm (green) and mitochondria (magenta).

**Figure S5.**
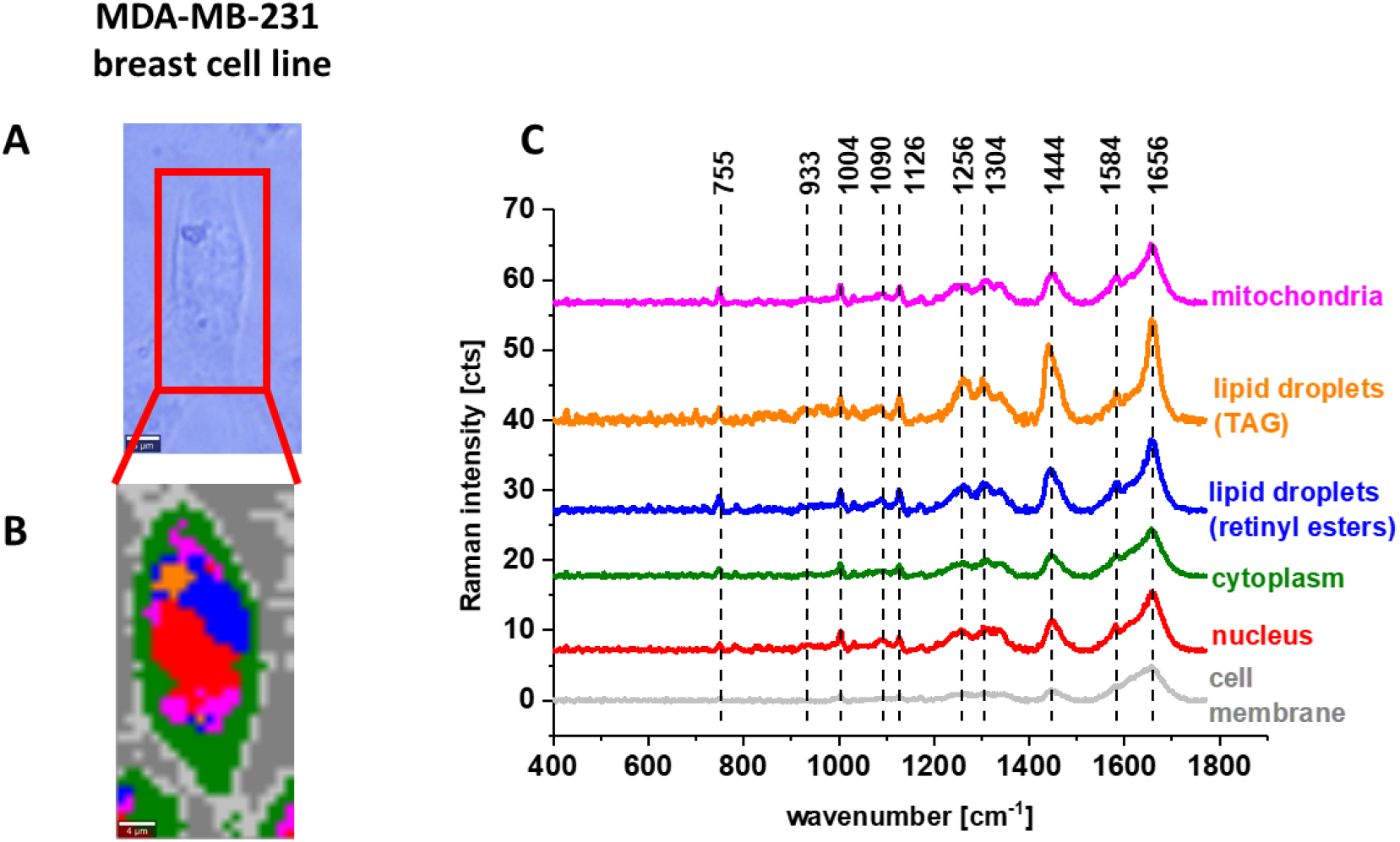
(A) The microscopy image, (B) the Raman image of a single cell MDA-MB-231 of highly aggressive breast cancer obtained by Cluster Analysis and (C) the corresponding average Raman spectra of different organelles of the cell: mitochondria (magenta), lipid droplets (TAG) (orange), lipid droplets (retinyl esters) (blue), cytoplasm (green), nucleus (red), cell membrane (light grey).

Figure S6 shows the intensity of Raman peak (raw data) centered at 1584 cm^-1^ as a function of Cyt c concentration for oxidized and reduced forms.

**Figure S6.**
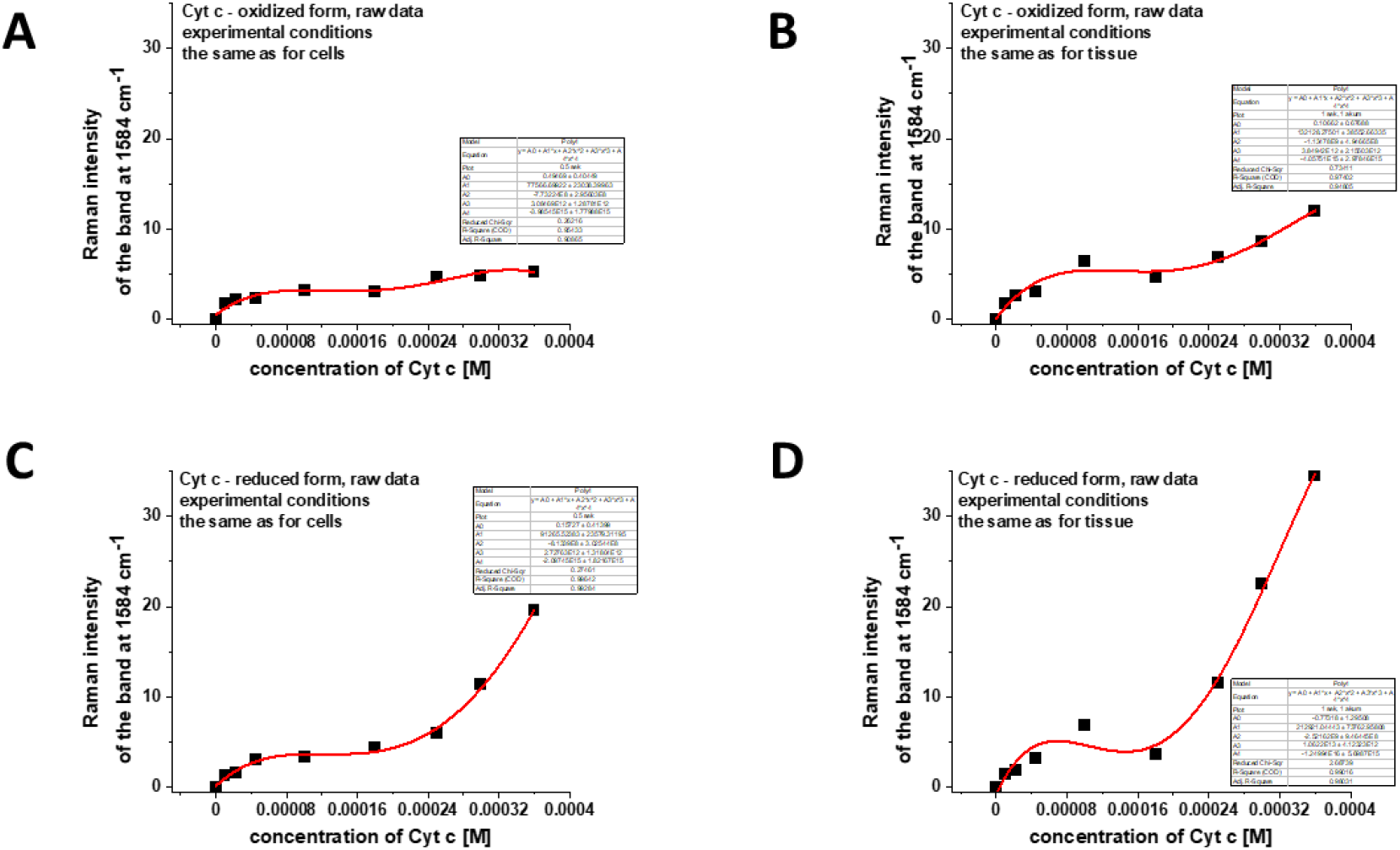
The intensity of Raman peak centered at 1584 cm^−1^ as a function of Cyt C concentration (raw data) for: (A) the oxidized form of Cyt c, experimental conditions the same as for breast single cells measurements: integration time 0.5 sec, 1 accumulation, laser power 10 mW; (B) the oxidized form of Cyt c, experimental conditions the same as for breast tissue measurements: integration time 1.0 sec, 1 accumulation, laser power 10 mW; (C) the reduced form of Cyt c, experimental conditions the same as for breast single cells measurements: integration time 0.5 sec, 1 accumulation, laser power 10 mW; (D) the reduced form of Cyt c, experimental conditions the same as for breast tissue measurements: integration time 1.0 sec, 1 accumulation, laser power 10 mW.

